# Spatiotemporal dynamics of EEG microstate networks over the first two years of life: A multi-cohort longitudinal study

**DOI:** 10.1101/2024.12.23.630142

**Authors:** Priyanka Ghosh, Kirsty Donald, Guilherme V. Polanczyk, Josh Paul Rodriguez, The Germina Project Team, The Khula Project Team, The LEAP 1kD EEG Working Group, Elizabeth Shephard, Laurel Gabard-Durnam

**Affiliations:** Center for Cognitive and Brain Health, Northeastern University, Boston, Massachusetts, USA; Division of Developmental Paediatrics, Department of Paediatrics and Child Health, Red Cross War Memorial Children’s Hospital, University of Cape Town, Cape Town, South Africa; The Neuroscience Institute, University of Cape Town, Cape Town, South Africa; Department of Psychiatry, School of Medicine, University of São Paulo, São Paulo, Brazil; Laboratory of Psychopathology and Psychiatric Therapeutics LIM-23, Institute of Psychiatry, Clinical Hospital HCFMUSP, School of Medicine, University of São Paulo, São Paulo, Brazil; Department of Developmental Learning and Personality Psychology, Institute of Psychology, University of São Paulo, São Paulo, Brazil

## Abstract

The first two years of life are marked by rapid development of large-scale brain networks that support emerging cognition and behavior. Magnetic-resonance approaches have revealed much about largescale networks in sleep, but very little is known about functional network dynamics in awake, behaving infants during this period of substantial development. Microstates are brief instances of distinct spatial topographies of largescale neural activity measured with electroencephalography (EEG) that offer a novel approach to studying whole-brain network dynamics at sub-second scale in awake infants by capturing their temporally coherent brain activity. While emerging literature is leveraging microstate dynamics in adults to understand mature largescale network function, developmental trajectories during networks’ rapid construction in infancy remain uncharacterized. In this study, we leveraged longitudinal resting-state EEG data from 854 infants across two geoculturally diverse cohorts to explore largescale network development through EEG microstates over the first two years of life. We provide evidence for conserved emergence of various network configurations (microstate classes A-G) through infancy across cohorts using data-driven clustering analyses. We also demonstrate significant longitudinal changes in microstate dynamics during this period, characterized by more numerous and more rapid transitions between largescale configurations, especially over early infancy. While patterns of sensory microstate development were largely consistent between cohorts, higher-order cognitive microstates showed context-specific developmental trends. Together these results provide novel insights into how large-scale brain networks functionally develop and organize across the first two years of life.

## Introduction

The first years of life represent a period of rapid and substantial brain development, including the first stages of functional maturation in largescale brain networks (Fair et al., 2009; Gabard-Durnam & McLaughlin, 2020; Paterson et al., 2006; Uddin et al., 2023). Large scale brain networks underlie interactions across widely distributed brain regions, supporting a range of sensory, cognitive, and socioemotional functions through coordinated neural activity (Bressler & Menon, 2010; Holz et al., 2023; Supekar et al., 2009). Most research on large-scale brain network development during the first years of life relies on functional magnetic resonance imaging (fMRI), which is limited by slower estimates of network dynamics, capturing changes on the scale of seconds to minutes, typically during sleep or sedation (Graham et al., 2015; Grayson & Fair, 2017; Smyser et al., 2011; Zhang et al., 2019). Moreover, resting state estimates of largescale networks often represent the average connectivity over recording periods of minutes, effectively treating network activity and between-network interactions as static (Custo et al., 2017; Hutchison et al., 2013; Zalesky et al., 2014). However, recent approaches suggest dynamic interactions between largescale networks unfold rapidly on the scale of milliseconds to seconds (Allen et al., 2012; Khazaei et al., 2023; Laumann et al., 2024). Consequently, there is currently extremely limited understanding of the spatiotemporal dynamics of largescale networks in awake, behaving infants and toddlers during this important period of brain and behavioral development (Desowska et al., 2024; Wang et al., 2021; Yates et al., 2023).

EEG’s temporal precision in indexing neural activity and suitability for awake recording from birth facilitates addressing this important gap in understanding the functional organization of the dynamic brain in early life. Specifically, EEG microstates capture instantaneous global brain network configurations and their dynamics (Allen et al., 2012; Michel & Koenig, 2018; Zanesco, 2024). EEG microstates are brief (60-120 ms in adults) periods of quasi-stable spatial configurations of neural activity shown to have close correspondence to the fMRI-derived largescale brain networks (Khanna et al., 2015; Tarailis et al., 2021; Lehmann et al., 1987; Michel & Koenig, 2018; Hill et al., 2023). Microstates measured during resting-state EEG in adults have distinct spatial (polarity-invariant) configurations associated with largescale brain networks typically labeled as A (right anterior to left posterior, associated with the auditory network), B (left anterior to right posterior; visual network), C (frontal to occipital; Default Mode Network), and D (medial anterior to occipital; Dorsal Attention Network) (Britz et al., 2010; Koenig et al., 2002, 2024;Khanna et al., 2014; Michel & Koenig, 2018). Though most studies primarily select these four canonical microstate configurations or classes apriori, more recent studies using data-driven approaches identify additional microstates and corresponding largescale networks, including microstate maps E (centro-parietal maximum, associated with Salience Network), F (left-lateralized maximum; anterior DMN) and G (right-lateralized maximum; sensorimotor network) (Custo et al., 2017; Das et al., 2022; Tarailis et al., 2023). The temporal pattern of resting state EEG microstates, including how frequently a given microstate is energetically-dominant and the average duration of dominance, indicates how largescale network configurations change and interact dynamically, offering insights into brain function of sensory, cognitive, and motor processes in both health and disease (Nazare & Tomescu, 2024; Niu et al., 2024; Tomescu et al., 2015;Hao et al., 2022; Khanna et al., 2015; Metzger et al., 2024).

While extensive research with EEG microstates in adults has revealed important information about largescale brain network dynamics and their relation to behavior, the field has only recently begun examining microstate indices of largescale network development in early life (Bagdasarov et al., 2024a; Bagdasarov et al., 2024b, Brown & Gartstein, 2023; Gui et al., 2021; Koenig et al. 2002; Khazaei et al., 2021; Hill et al., 2023). Preliminary studies measuring microstates cross-sectionally indicate that EEG microstates in early life may exhibit different characteristics compared to adults and offer important insight into early development (Bagdasarov et al., 2022; Hill et al., 2023; Koenig et al., 2002). For example, Brown and Gartstein (2023) have shown that microstates extracted from 6- to 10-month olds have concurrent associations with individual differences in temperament, suggesting early behavioral significance of infant microstate dynamics. Gui et al. (2021) have shown that infant microstate dynamics are also indicative of behavior measured years later where microstates during a social attention paradigm at 8 months of age were related to subsequent social skills and Autism Spectrum Disorder diagnoses at 3 years of age. These initial findings with cross-sectional microstate measurements establish that infant microstate dynamics are important for understanding largescale brain networks and emerging behavior. However, the normative longitudinal development of microstate configurations and their dynamics over infancy remain unknown.

Our study therefore aims to shed light on the development of largescale brain network dynamics over infancy by analyzing the spatiotemporal dynamics of resting state EEG microstates longitudinally over the first two years of life. We conducted a multi-site longitudinal study with harmonized EEG acquisitions across two cohorts (n=854) that differ in their geographic, cultural, demographic, and socioeconomic characteristics drawn from Cape Town, South Africa, and São Paulo, Brazil. We characterized the developmental trajectory of EEG microstates in each sample of infants longitudinally as a function of age and sex and compared their developmental patterns to examine the consistency of developmental changes across contexts. By examining how EEG microstates evolve over infancy, this study seeks to improve understanding of early largescale brain network dynamics when they rapidly develop and to provide insights into the neural underpinnings of early sensory, cognitive, and behavioral development.

## Results

### Developmental changes in microstate classes

As shown in **Figure 1**, polarity-invariant canonical microstates A, B, C, D and E were seen in individuals at 2-6 months as well as at 5-12 months in the Khula cohort. Microstate C was replaced with the emergence of two additional microstates F and G at 12-18 months and 19-26 months in the Khula cohort. In the Germina cohort, microstates A, B, C, D and E were seen at 3-4 months, 5-10 months and 10-17 months. Microstate C was replaced with the emergence of microstates F and G at 18-30 months. Microstate statistics for all classes (A through G) at each timepoint for each cohort are depicted in **Table 3**.

**Figure 1.**
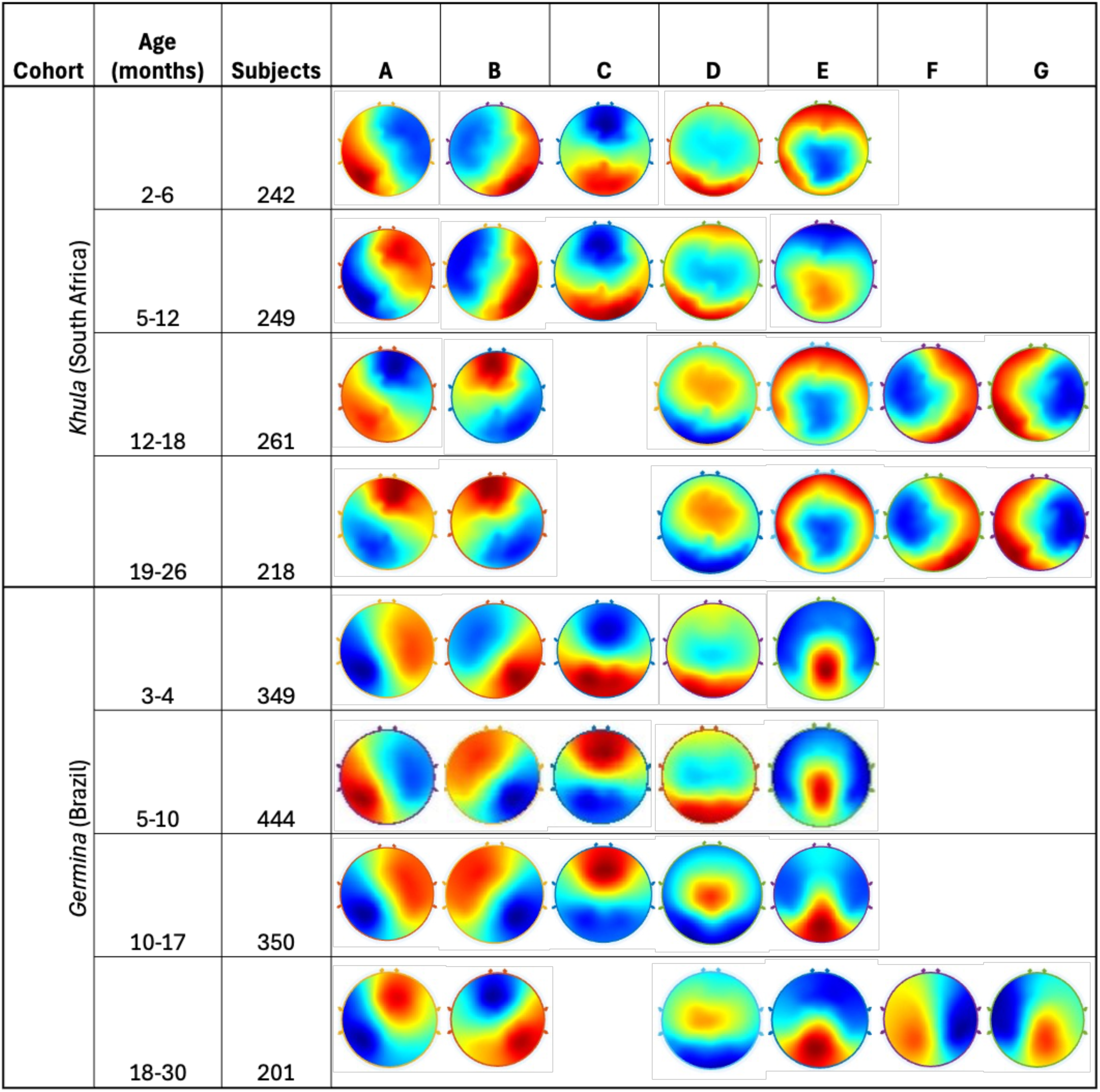
Infant microstates. The figure represents open-eyes resting state EEG microstate topographies of infants recorded at four different developmental time windows longitudinally across the first two years of life from South Africa based *Khula* and Brazil based *Germina* cohorts.

**Figure 2.**
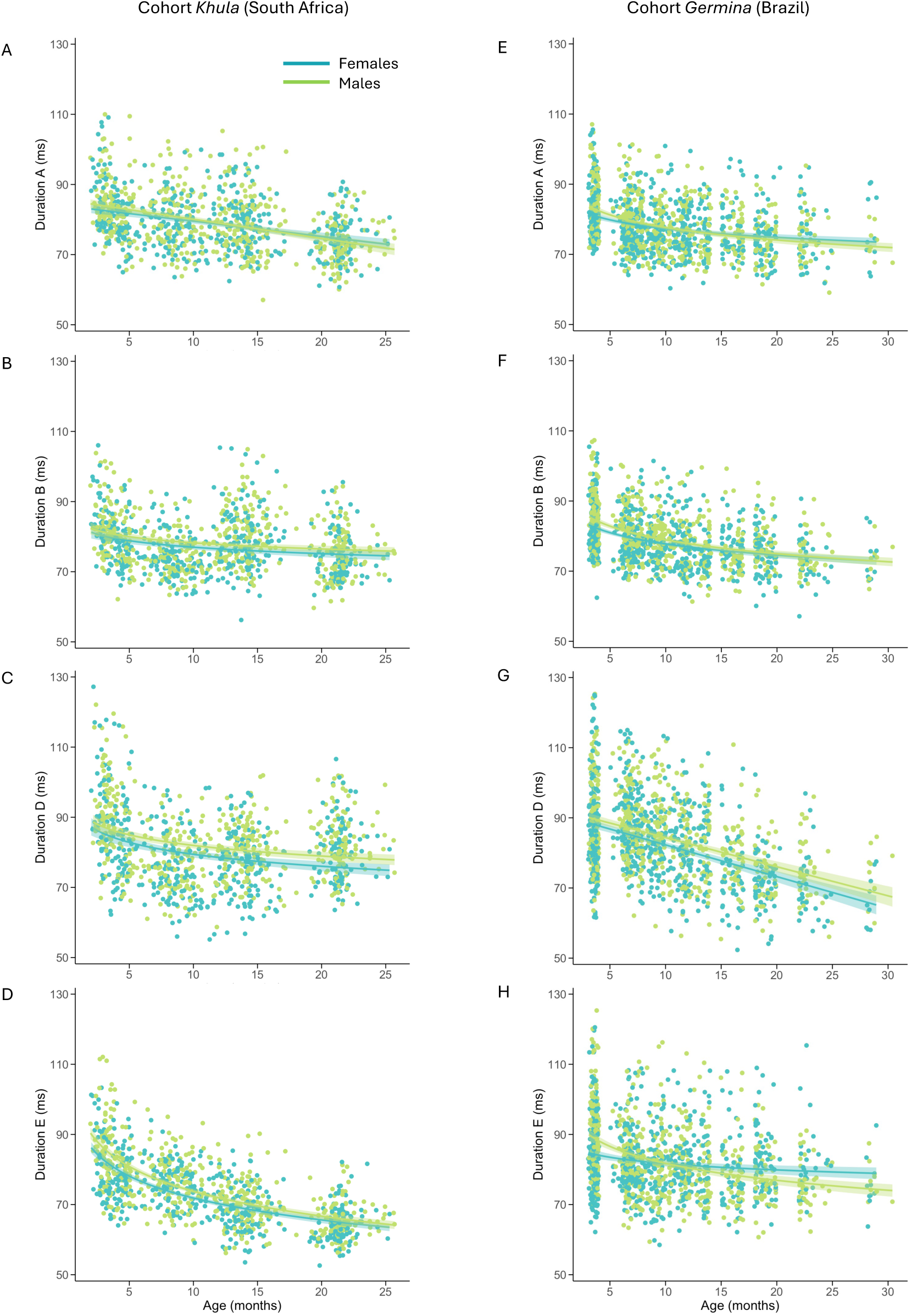
Developmental trajectories of microstate durations through infancy. The figure shows cohort specific age-associated changes in the duration of microstates A(2A,2E), B(2B,2F), D(3C,3G) and E(3D,3H), categorized by sex (light green=males; teal=females). Linear or logarithmic regression lines and their 95% confidence intervals (shaded) are shown in each plot.

**Figure 3.**
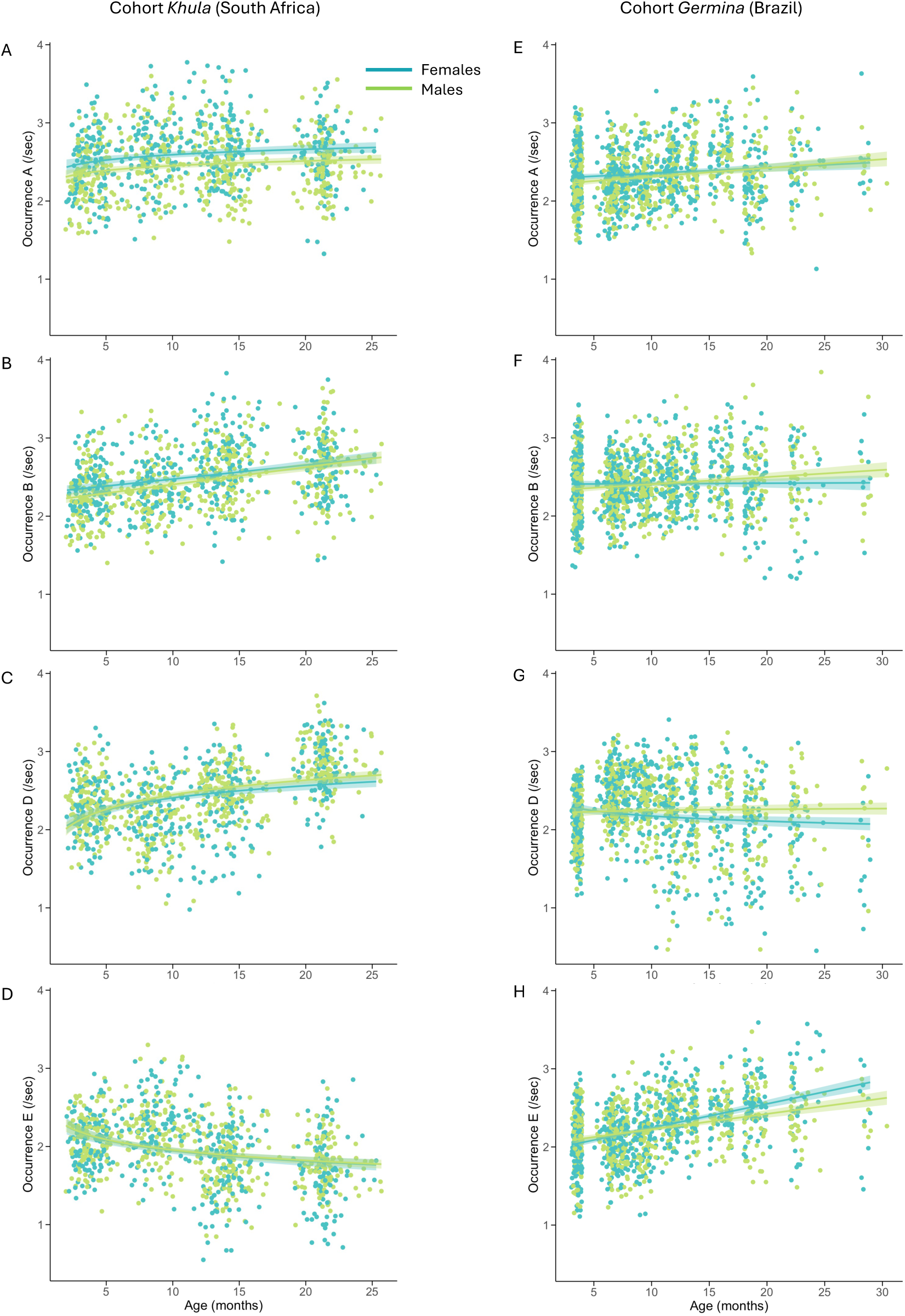
Developmental trajectories of microstate occurrences through infancy. The figure shows cohort specific age-associated changes in the occurrence of microstates A(2A,2E), B(2B,2F), D(3C,3G) and E(3D,3H), categorized by sex (light green=males; teal=females). Linear or logarithmic regression lines and their 95% confidence intervals (shaded) are shown in each plot.

**Table 1.**
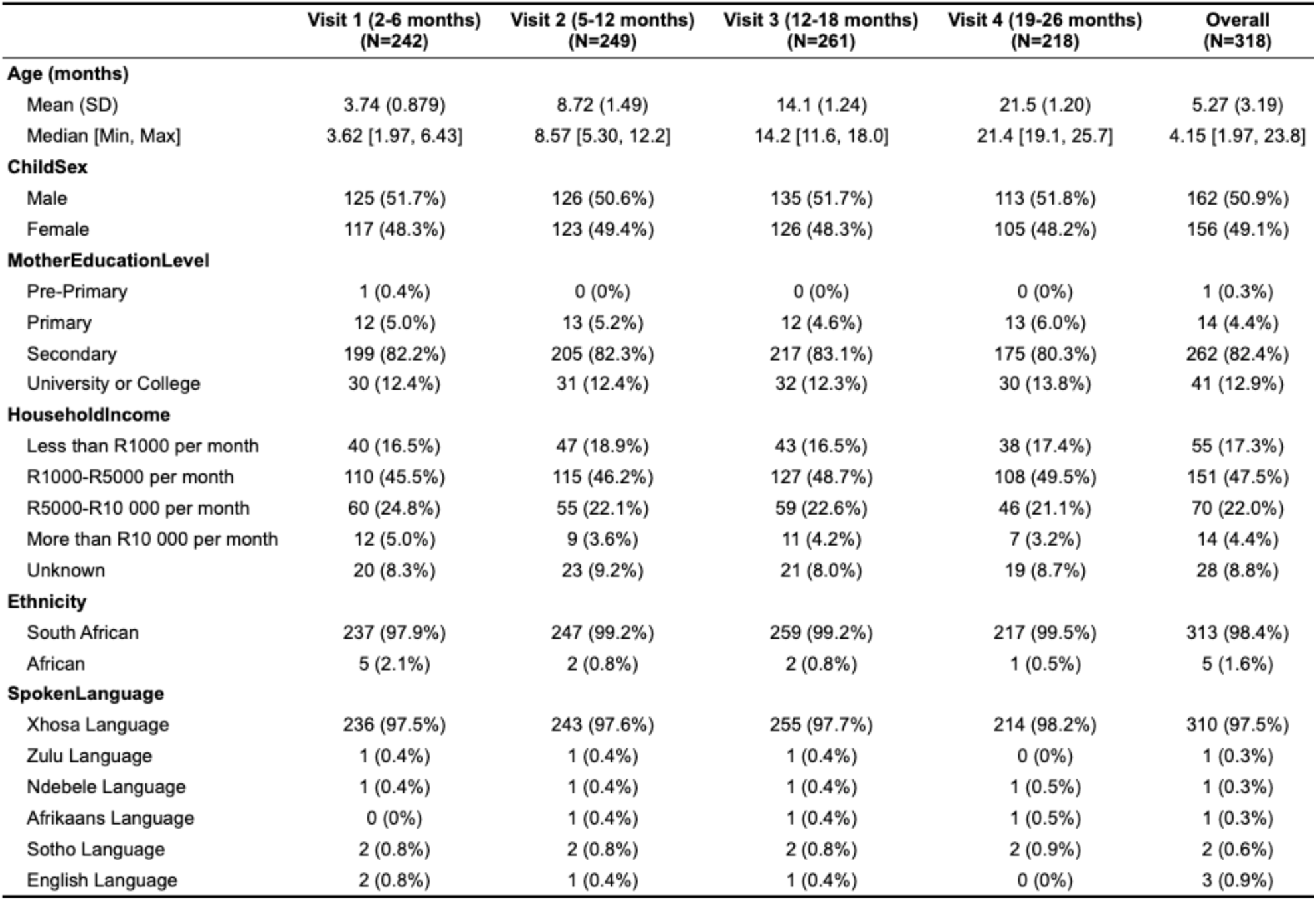
Descriptive statistics of demographic data from the South Africa based *Khula* study.

**Table 2.** Descriptive statistics of demographic data from the Brazil based *Germina* study.

|  | Visit 1 (3-4 months)<br>(N=349) | Visit 2 (5-10 months)<br>(N=444) | Visit 3 (10-17 months)<br>(N=350) | Visit 4 (18-30 months)<br>(N=201) | Overall<br>(N=536) |
| --- | --- | --- | --- | --- | --- |
| **Age (months)** |  |  |  |  |  |
| Mean (SD) | 3.62 (0.213) | 7.64 (1.25) | 13.2 (2.06) | 21.1 (3.11) | 5.41 (3.27) |
| Median [Min, Max] | 3.61 [3.06, 4.01] | 7.41 [5.06, 9.99] | 12.9 [10.0, 17.0] | 19.7 [18.0, 30.4] | 3.81 [3.06, 28.8] |
| **ChildSex** |  |  |  |  |  |
| Male | 158 (45.3%) | 217 (48.9%) | 171 (48.9%) | 101 (50.2%) | 261 (48.7%) |
| Female | 191 (54.7%) | 227 (51.1%) | 179 (51.1%) | 100 (49.8%) | 275 (51.3%) |
| **MotherEducationLevel** |  |  |  |  |  |
| Primary | 5 (1.4%) | 4 (0.9%) | 3 (0.9%) | 2 (1.0%) | 5 (0.9%) |
| Secondary | 71 (20.3%) | 96 (21.6%) | 76 (21.7%) | 40 (19.9%) | 108 (20.1%) |
| University or College | 273 (78.2%) | 344 (77.5%) | 271 (77.4%) | 159 (79.1%) | 423 (78.9%) |
| **FatherEducationLevel** |  |  |  |  |  |
| Primary | 17 (4.9%) | 17 (3.8%) | 14 (4.0%) | 6 (3.0%) | 23 (4.3%) |
| Secondary | 87 (24.9%) | 112 (25.2%) | 87 (24.9%) | 61 (30.3%) | 131 (24.4%) |
| University or College | 241 (69.1%) | 310 (69.8%) | 246 (70.3%) | 131 (65.2%) | 377 (70.3%) |
| Unknown | 4 (1.1%) | 5 (1.1%) | 3 (0.9%) | 3 (1.5%) | 5 (0.9%) |
| **HouseholdIncome** |  |  |  |  |  |
| Less than BRL1000 per month | 19 (5.4%) | 10 (2.3%) | 3 (0.9%) | 1 (0.5%) | 20 (3.7%) |
| BRL1000-BRL5000 per month | 131 (37.5%) | 129 (29.1%) | 97 (27.7%) | 69 (34.3%) | 182 (34.0%) |
| BRL5000-BRL10 000 per month | 80 (22.9%) | 118 (26.6%) | 107 (30.6%) | 51 (25.4%) | 136 (25.4%) |
| More than BRL10 000 per month | 118 (33.8%) | 160 (36.0%) | 126 (36.0%) | 66 (32.8%) | 183 (34.1%) |
| Unknown | 1 (0.3%) | 27 (6.1%) | 17 (4.9%) | 14 (7.0%) | 15 (2.8%) |
| **Ethnicity** |  |  |  |  |  |
| White | 225 (64.5%) | 286 (64.4%) | 224 (64.0%) | 129 (64.2%) | 349 (65.1%) |
| Black | 34 (9.7%) | 40 (9.0%) | 28 (8.0%) | 18 (9.0%) | 46 (8.6%) |
| Pardo | 72 (20.6%) | 92 (20.7%) | 74 (21.1%) | 42 (20.9%) | 111 (20.7%) |
| Other | 18 (5.2%) | 26 (5.9%) | 24 (6.9%) | 12 (6.0%) | 30 (5.6%) |
| **SpokenLanguage** |  |  |  |  |  |
| Portuguese Language | 344 (98.6%) | 439 (98.9%) | 347 (99.1%) | 199 (99.0%) | 529 (98.7%) |
| Spanish Language | 2 (0.6%) | 2 (0.5%) | 1 (0.3%) | 0 (0%) | 2 (0.4%) |
| Bilingual | 1 (0.3%) | 0 (0%) | 0 (0%) | 0 (0%) | 1 (0.2%) |
| English Language | 0 (0%) | 1 (0.2%) | 1 (0.3%) | 1 (0.5%) | 2 (0.4%) |
| Unknown | 2 (0.6%) | 2 (0.5%) | 1 (0.3%) | 1 (0.5%) | 2 (0.4%) |

**Table 3.**
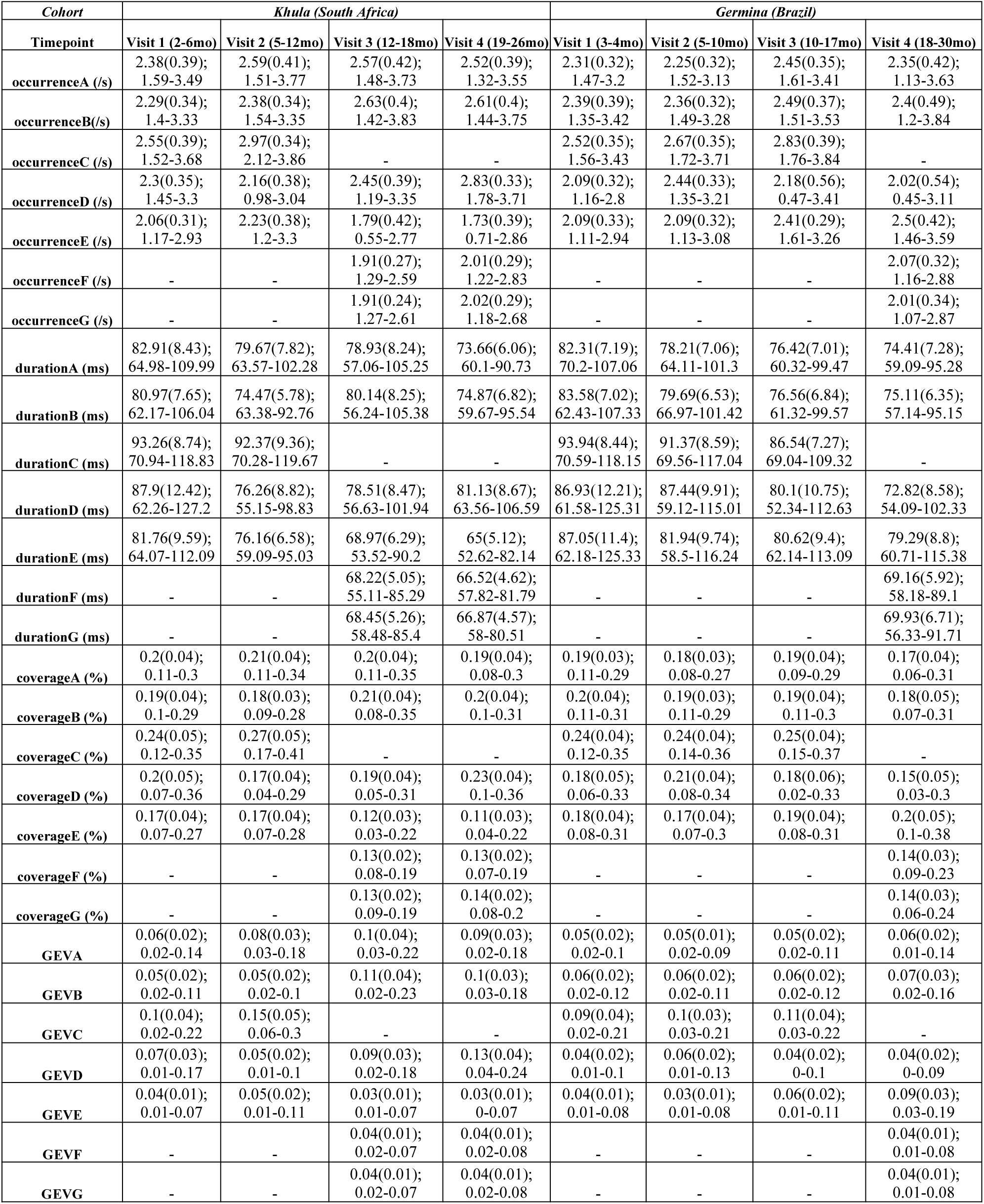
Summary statistics of microstate parameters. Each cell in the table lists the mean, standard deviation, minimum and maximum values (mean(SD); min-max) of the following microstate temporal parameters: Occurrence, Duration, Coverage and Global Explained Variance (GEV) of classes A, B, C, D, E, F and G for participants from the *Khula* and *Germina* cohorts for Visit 1 through 4.

| *Cohort* | *Khula (South Africa)* |  |  |  | *Germina (Brazil)* |  |  |  |
| --- | --- | --- | --- | --- | --- | --- | --- | --- |
| **Timepoint** | **Visit 1 (2-6mo)** | **Visit 2 (5-12mo)** | **Visit 3 (12-18mo)** | **Visit 4 (19-26mo)** | **Visit 1 (3-4mo)** | **Visit 2 (5-10mo)** | **Visit 3 (10-17mo)** | **Visit 4 (18-30mo)** |
| **occurrenceA (/s)** | 2.38(0.39);<br>1.59-3.49 | 2.59(0.41);<br>1.51-3.77 | 2.57(0.42);<br>1.48-3.73 | 2.52(0.39);<br>1.32-3.55 | 2.31(0.32);<br>1.47-3.2 | 2.25(0.32);<br>1.52-3.13 | 2.45(0.35);<br>1.61-3.41 | 2.35(0.42);<br>1.13-3.63 |
| **occurrenceB(/s)** | 2.29(0.34);<br>1.4-3.33 | 2.38(0.34);<br>1.54-3.35 | 2.63(0.4);<br>1.42-3.83 | 2.61(0.4);<br>1.44-3.75 | 2.39(0.39);<br>1.35-3.42 | 2.36(0.32);<br>1.49-3.28 | 2.49(0.37);<br>1.51-3.53 | 2.4(0.49);<br>1.2-3.84 |
| **occurrenceC (/s)** | 2.55(0.39);<br>1.52-3.68 | 2.97(0.34);<br>2.12-3.86 | - | - | 2.52(0.35);<br>1.56-3.43 | 2.67(0.35);<br>1.72-3.71 | 2.83(0.39);<br>1.76-3.84 | - |
| **occurrenceD (/s)** | 2.3(0.35);<br>1.45-3.3 | 2.16(0.38);<br>0.98-3.04 | 2.45(0.39);<br>1.19-3.35 | 2.83(0.33);<br>1.78-3.71 | 2.09(0.32);<br>1.16-2.8 | 2.44(0.33);<br>1.35-3.21 | 2.18(0.56);<br>0.47-3.41 | 2.02(0.54);<br>0.45-3.11 |
| **occurrenceE (/s)** | 2.06(0.31);<br>1.17-2.93 | 2.23(0.38);<br>1.2-3.3 | 1.79(0.42);<br>0.55-2.77 | 1.73(0.39);<br>0.71-2.86 | 2.09(0.33);<br>1.11-2.94 | 2.09(0.32);<br>1.13-3.08 | 2.41(0.29);<br>1.61-3.26 | 2.5(0.42);<br>1.46-3.59 |
| **occurrenceF (/s)** | - | - | 1.91(0.27);<br>1.29-2.59 | 2.01(0.29);<br>1.22-2.83 | - | - | - | 2.07(0.32);<br>1.16-2.88 |
| **occurrenceG (/s)** | - | - | 1.91(0.24);<br>1.27-2.61 | 2.02(0.29);<br>1.18-2.68 | - | - | - | 2.01(0.34);<br>1.07-2.87 |
| **durationA (ms)** | 82.91(8.43);<br>64.98-109.99 | 79.67(7.82);<br>63.57-102.28 | 78.93(8.24);<br>57.06-105.25 | 73.66(6.06);<br>60.1-90.73 | 82.31(7.19);<br>70.2-107.06 | 78.21(7.06);<br>64.11-101.3 | 76.42(7.01);<br>60.32-99.47 | 74.41(7.28);<br>59.09-95.28 |
| **durationB (ms)** | 80.97(7.65);<br>62.17-106.04 | 74.47(5.78);<br>63.38-92.76 | 80.14(8.25);<br>56.24-105.38 | 74.87(6.82);<br>59.67-95.54 | 83.58(7.02);<br>62.43-107.33 | 79.69(6.53);<br>66.97-101.42 | 76.56(6.84);<br>61.32-99.57 | 75.11(6.35);<br>57.14-95.15 |
| **durationC (ms)** | 93.26(8.74);<br>70.94-118.83 | 92.37(9.36);<br>70.28-119.67 | - | - | 93.94(8.44);<br>70.59-118.15 | 91.37(8.59);<br>69.56-117.04 | 86.54(7.27);<br>69.04-109.32 | - |
| **durationD (ms)** | 87.9(12.42);<br>62.26-127.2 | 76.26(8.82);<br>55.15-98.83 | 78.51(8.47);<br>56.63-101.94 | 81.13(8.67);<br>63.56-106.59 | 86.93(12.21);<br>61.58-125.31 | 87.44(9.91);<br>59.12-115.01 | 80.1(10.75);<br>52.34-112.63 | 72.82(8.58);<br>54.09-102.33 |
| **durationE (ms)** | 81.76(9.59);<br>64.07-112.09 | 76.16(6.58);<br>59.09-95.03 | 68.97(6.29);<br>53.52-90.2 | 65(5.12);<br>52.62-82.14 | 87.05(11.4);<br>62.18-125.33 | 81.94(9.74);<br>58.5-116.24 | 80.62(9.4);<br>62.14-113.09 | 79.29(8.8);<br>60.71-115.38 |
| **durationF (ms)** | - | - | 68.22(5.05);<br>55.11-85.29 | 66.52(4.62);<br>57.82-81.79 | - | - | - | 69.16(5.92);<br>58.18-89.1 |
| **durationG (ms)** | - | - | 68.45(5.26);<br>58.48-85.4 | 66.87(4.57);<br>58-80.51 | - | - | - | 69.93(6.71);<br>56.33-91.71 |
| **coverageA (%)** | 0.2(0.04);<br>0.11-0.3 | 0.21(0.04);<br>0.11-0.34 | 0.2(0.04);<br>0.11-0.35 | 0.19(0.04);<br>0.08-0.3 | 0.19(0.03);<br>0.11-0.29 | 0.18(0.03);<br>0.08-0.27 | 0.19(0.04);<br>0.09-0.29 | 0.17(0.04);<br>0.06-0.31 |
| **coverageB (%)** | 0.19(0.04);<br>0.1-0.29 | 0.18(0.03);<br>0.09-0.28 | 0.21(0.04);<br>0.08-0.35 | 0.2(0.04);<br>0.1-0.31 | 0.2(0.04);<br>0.11-0.31 | 0.19(0.03);<br>0.11-0.29 | 0.19(0.04);<br>0.11-0.3 | 0.18(0.05);<br>0.07-0.31 |
| **coverageC (%)** | 0.24(0.05);<br>0.12-0.35 | 0.27(0.05);<br>0.17-0.41 | - | - | 0.24(0.04);<br>0.12-0.35 | 0.24(0.04);<br>0.14-0.36 | 0.25(0.04);<br>0.15-0.37 | - |
| **coverageD (%)** | 0.2(0.05);<br>0.07-0.36 | 0.17(0.04);<br>0.04-0.29 | 0.19(0.04);<br>0.05-0.31 | 0.23(0.04);<br>0.1-0.36 | 0.18(0.05);<br>0.06-0.33 | 0.21(0.04);<br>0.08-0.34 | 0.18(0.06);<br>0.02-0.33 | 0.15(0.05);<br>0.03-0.3 |
| **coverageE (%)** | 0.17(0.04);<br>0.07-0.27 | 0.17(0.04);<br>0.07-0.28 | 0.12(0.03);<br>0.03-0.22 | 0.11(0.03);<br>0.04-0.22 | 0.18(0.04);<br>0.08-0.31 | 0.17(0.04);<br>0.07-0.3 | 0.19(0.04);<br>0.08-0.31 | 0.2(0.05);<br>0.1-0.38 |
| **coverageF (%)** | - | - | 0.13(0.02);<br>0.08-0.19 | 0.13(0.02);<br>0.07-0.19 | - | - | - | 0.14(0.03);<br>0.09-0.23 |
| **coverageG (%)** | - | - | 0.13(0.02);<br>0.09-0.19 | 0.14(0.02);<br>0.08-0.2 | - | - | - | 0.14(0.03);<br>0.06-0.24 |
| **GEVA** | 0.06(0.02);<br>0.02-0.14 | 0.08(0.03);<br>0.03-0.18 | 0.1(0.04);<br>0.03-0.22 | 0.09(0.03);<br>0.02-0.18 | 0.05(0.02);<br>0.02-0.1 | 0.05(0.01);<br>0.02-0.09 | 0.05(0.02);<br>0.02-0.11 | 0.06(0.02);<br>0.01-0.14 |
| **GEVB** | 0.05(0.02);<br>0.02-0.11 | 0.05(0.02);<br>0.02-0.1 | 0.11(0.04);<br>0.02-0.23 | 0.1(0.03);<br>0.03-0.18 | 0.06(0.02);<br>0.02-0.12 | 0.06(0.02);<br>0.02-0.11 | 0.06(0.02);<br>0.02-0.12 | 0.07(0.03);<br>0.02-0.16 |
| **GEVC** | 0.1(0.04);<br>0.02-0.22 | 0.15(0.05);<br>0.06-0.3 | - | - | 0.09(0.04);<br>0.02-0.21 | 0.1(0.03);<br>0.03-0.21 | 0.11(0.04);<br>0.03-0.22 | - |
| **GEVD** | 0.07(0.03);<br>0.01-0.17 | 0.05(0.02);<br>0.01-0.1 | 0.09(0.03);<br>0.02-0.18 | 0.13(0.04);<br>0.04-0.24 | 0.04(0.02);<br>0.01-0.1 | 0.06(0.02);<br>0.01-0.13 | 0.04(0.02);<br>0-0.1 | 0.04(0.02);<br>0-0.09 |
| **GEVE** | 0.04(0.01);<br>0.01-0.07 | 0.05(0.02);<br>0.01-0.11 | 0.03(0.01);<br>0.01-0.07 | 0.03(0.01);<br>0-0.07 | 0.04(0.01);<br>0.01-0.08 | 0.03(0.01);<br>0.01-0.08 | 0.06(0.02);<br>0.01-0.11 | 0.09(0.03);<br>0.03-0.19 |
| **GEVF** | - | - | 0.04(0.01);<br>0.02-0.07 | 0.04(0.01);<br>0.02-0.08 | - | - | - | 0.04(0.01);<br>0.01-0.08 |
| **GEVG** | - | - | 0.04(0.01);<br>0.02-0.07 | 0.04(0.01);<br>0.02-0.08 | - | - | - | 0.04(0.01);<br>0.01-0.08 |

### Developmental changes in microstate dynamics

To assess developmental changes in microstate dynamics over the first 2 years of life, primary analyses focused on changes in microstate occurrences (i.e., average number of times a given microstate was present per second) and durations (i.e., average time in milliseconds that a given microstate was uninterruptedly present within the EEG) over this age range. Specifically, occurrence and duration features were assessed for the microstate classes present across all four timepoints (i.e., microstates A, B, D and E) using linear mixed-effects models. Models included number of EEG segments retained as covariates of no interest in addition to effects of sex and age (linear). Additional non-linear age-related changes were tested for via inclusion of added nonlinear terms (quadratic or logarithmic) and retained if either non-linear term improved model fit. We observed developmental changes in microstates’ duration and occurrence statistics (**Tables 4 and 5**) which are detailed for each microstate below.

**Table 4.** Developmental changes in microstate durations. The statistics for fixed effects of the model outputs (durationA/B/D/E) have been listed as estimates and their standard errors for microstate durations being predicted by age and sex of infants, and the number of segments retained from each EEG recording. The effect size for the significant regressions (p<0.05) were estimated using partial eta squared (η_p_ ^2^) categorized as small (denoted by *) if η_p_ ^2^20.01, medium (denoted by **) if η_p_ ^2^20.06 or large (denoted by ***) if η_p_ ^2^20.14 based on Cohen, 2013.

| Cohort | Microstate | Model (age) | Predictor | Estimate (b) | Std. Error | p-value | Effect size($\eta_p^2$) |
| --- | --- | --- | --- | --- | --- | --- | --- |
| *Khula* | durationA | linear | age | -0.505 | 0.036 | **<0.001** | ***0.200 |
|  |  |  | sex | 0.305 | 0.582 | 0.600 | 0.001 |
|  |  |  | segments | -0.042 | 0.038 | 0.268 | 0.001 |
|  | durationB | linear + log | age | 0.364 | 0.131 | **0.006** | *0.009 |
|  |  |  | log(age) | -5.925 | 1.283 | **<0.001** | *0.025 |
|  |  |  | sex | 1.172 | 0.504 | **0.021** | *0.019 |
|  |  |  | segments | 0.012 | 0.011 | 0.265 | 0.001 |
|  | durationD | linear + log | age | 1.902 | 0.153 | **<0.001** | ***0.170 |
|  |  |  | log(age) | -22.407 | 1.502 | **<0.001** | ***0.224 |
|  |  |  | sex | 2.461 | 0.745 | **0.001** | *0.035 |
|  |  |  | segments | 0.000 | 0.013 | 0.982 | 0.000 |
|  | durationE | linear + log | age | -0.259 | 0.113 | **0.022** | *0.007 |
|  |  |  | log(age) | -7.160 | 1.108 | **<0.001** | *0.052 |
|  |  |  | sex | 1.944 | 0.540 | **<0.001** | *0.043 |
|  |  |  | segments | 0.027 | 0.010 | 0.006 | 0.009 |
| *Germina* | durationA | linear + log | age | 0.234 | 0.106 | **0.027** | *0.005 |
|  |  |  | log(age) | -6.575 | 1.031 | **<0.001** | *0.038 |
|  |  |  | sex | 0.654 | 0.440 | 0.137 | 0.004 |
|  |  |  | segments | -0.029 | 0.017 | 0.083 | 0.002 |
|  | durationB | linear + log | age | 0.094 | 0.101 | 0.350 | 0.001 |
|  |  |  | log(age) | -5.958 | 0.981 | **<0.001** | *0.034 |
|  |  |  | sex | 1.274 | 0.400 | **0.002** | *0.019 |
|  |  |  | segments | 0.032 | 0.016 | 0.043 | 0.003 |
|  | durationD | linear | age | -0.880 | 0.047 | **<0.001** | ***0.230 |
|  |  |  | sex | 2.193 | 0.631 | **0.005** | *0.022 |
|  |  |  | segments | -0.034 | 0.025 | 0.180 | 0.001 |
|  | durationE | linear + log | age | 0.310 | 0.147 | **0.035** | 0.004 |
|  |  |  | log(age) | -7.507 | 1.430 | **<0.001** | *0.026 |
|  |  |  | sex | 0.970 | 0.621 | 0.119 | 0.005 |
|  |  |  | segments | -0.018 | 0.024 | 0.454 | 0.000 |

**Table 5.**
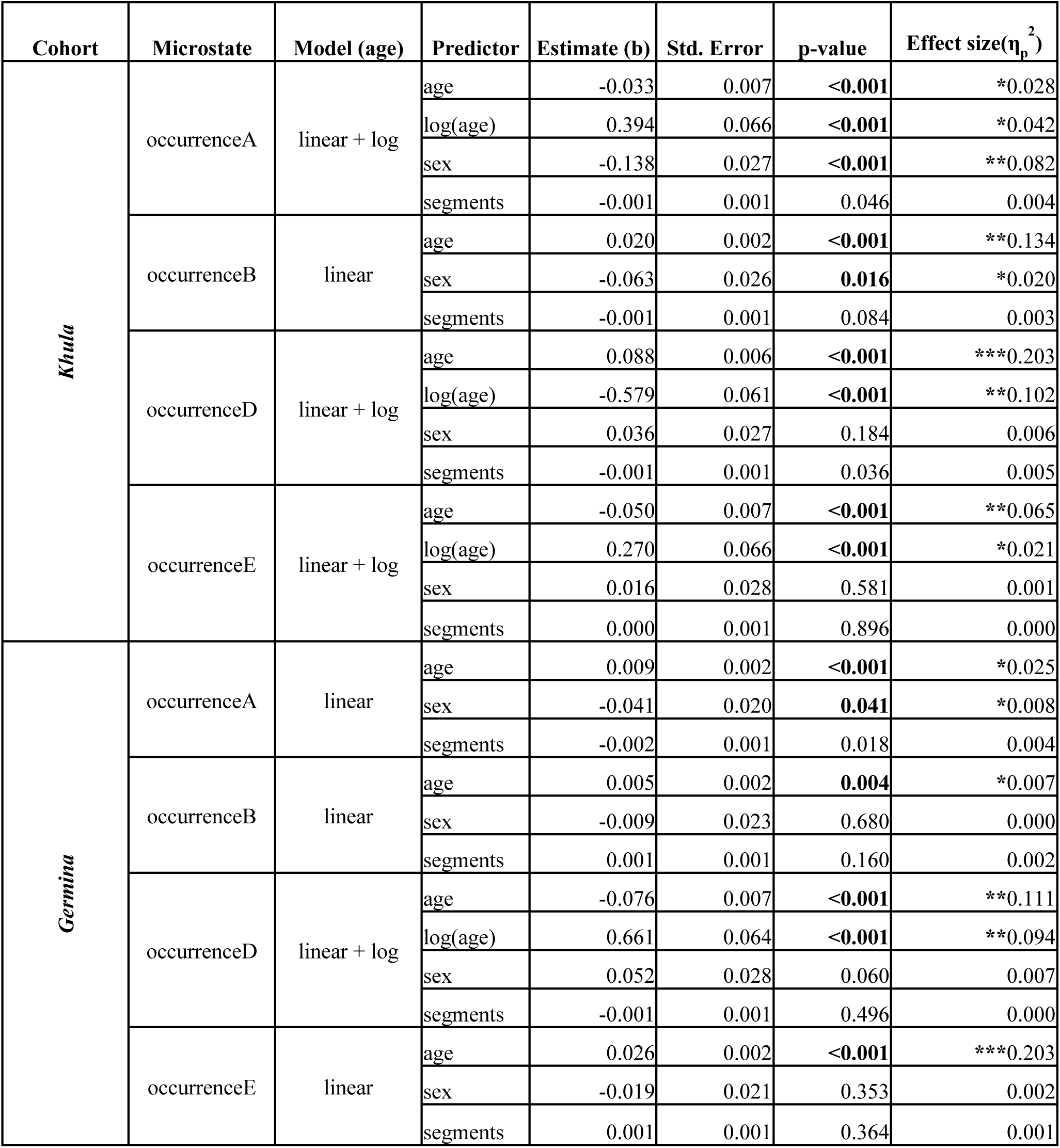
Developmental changes in microstate occurrences. The statistics for fixed effects of the model outputs (occurrenceA/B/D/E) have been listed as estimates and their standard errors for microstate occurrences being predicted by age and sex of infants, and the number of segments retained from each EEG recording. The effect size for the significant regressions (p<0.05) were estimated using partial eta squared (η_p_^2^) categorized as small (denoted by *) if η_p_^2^20.01, medium (denoted by **) if η_p_^2^20.06 or large (denoted by ***) if η_p_^2^20.14 based on Cohen (2013).

#### Microstate A

Microstate A durations significantly decreased with age across both cohorts over infancy. The linear mixed effects model in Khula (South Africa) showed a strong negative effect of linear age on microstate A duration (b=-0.505, p<0.001, η_p_^2^=0.2). The model in Germina (Brazil) revealed a non-linear negative effect of age on microstate A duration (b=-6.575, p<0.001, η_p_^2^=0.038), such that duration significantly decreased with increasing age, with faster rates of change in early infancy. Infant sex did not differentiate developmental trajectories for microstate A duration in either Khula or Germina (p > 0.05).

Though microstate A decreased in duration, it significantly increased its occurrence with age. This relation was non-linear within individuals in the Khula cohort (b=0.394, p<0.001, η_p_^2^=0.042, increasing most quickly over early infancy) and linear in the Germina cohort (b=0.009, p<0.001, η_p_^2^=0.025). Moreover, the infant’s biological sex was a significant predictor of the occurrence of microstate A in both Khula (b=-0.138, p<0.001, η_p_^2^=0.082) and Germina (b=-0.041, p=0.041, η_p_^2^=0.008), where males demonstrated fewer microstate A occurrences than females, engaging this largescale brain network less frequently each second.

#### Microstate B

Microstate B durations significantly decreased non-linearly with age across both cohorts over infancy. The Khula mixed effects model revealed a non-linear effect of age on microstate B duration within individuals, with duration decreasing most steeply over the first year (b=-5.925, p<0.001, η_p_^2^=0.025). Similar trends were seen in Germina (b=-5.958, p<0.001, η_p_^2^=0.034). Additionally, infant sex had a significant but small effect in predicting the duration of Microstate B in both Khula (b=1.172, p=0.021, η_p_^2^=0.019) and Germina (b=1.274, p=0.002, η_p_^2^=0.019), such that males had significantly longer microstate durations than females, indicating slower transitions from Microstate B to other microstate configurations in males.

Microstate B linearly increased in occurrence over the first 2 years in both cohorts (Khula: b=0.020, p<0.001, η_p_^2^=0.134; Germina: b=0.005, p=0.004, η_p_^2^=0.007). The occurrence of microstate B was also significantly predicted by infant sex in Khula (b=-0.063, p=0.016, η_p_^2^=0.020) with males demonstrating fewer occurrences than females, though this effect was small.

#### Microstate D

Microstate D durations significantly decreased with age with large developmental effects in both cohorts. The linear mixed effects model for Khula revealed a strong non-linear effect of age on microstate D duration within individuals, with duration showing the most rapid decrease in early infancy (b=-22.407, p<0.001, η_p_^2^=0.224). In Germina, a strong negative linear effect of age was observed on microstate D duration within individuals (b= −0.880, p<0.001, η_p_^2^=0.230). Additionally, infant sex had a significant effect in predicting the duration of Microstate D in both Khula (b=2.461, p=0.001, η_p_^2^=0.035) and Germina (b=2.193, p=0.005, η_p_^2^=0.022) such that males had significantly longer microstate durations than females, though this effect was small in both cohorts.

Microstate D occurrence demonstrated different non-linear developmental patterns across cohorts. In Khula, there was a strong non-linear increase in microstate D occurrence with age with the steepest increase in the second year (b=-0.579, p<0.001, η_p_^2^=0.102). However, the model for Germina suggested an early increase followed by a decrease over the second year in occurrence (b=0.661, p<0.001, η_p_^2^=0.094). Infant sex did not differentiate developmental trajectories for microstate D occurrence in either Khula or Germina (p > 0.05).

#### Microstate E

Microstate E decreased in duration non-linearly with age in both cohorts, with the fastest rate of change earlier in infancy (Khula: b=-7.160, p<0.001, η_p_^2^=0.052; Germina: b=-7.507, p<0.001, η_p_^2^=0.026). Moreover, infant sex in Khula significantly (b=1.944, p<0.001, η_p_^2^=0.043) predicted the duration of Microstate E, where males had longer microstate durations than females.

Microstate E occurrence demonstrated different non-linear developmental patterns across cohorts. The mixed effects model in Khula indicated a non-linear decrease in occurrence, most steeply around the end of the first year of life (b=0.270, p<0.001, η_p_^2^=0.021). In Germina, a strong, positive linear effect of age was observed on the occurrence of microstate E (b=0.026, p<0.001, η_p_^2^=0.203). For both Khula and Germina, infant sex did not differentiate developmental trajectories of microstate E occurrence (p > 0.05).

## Discussion

Our study provides a comprehensive characterization of resting state EEG microstate dynamics during the first two years of life when the most rapid and significant development of largescale functional networks takes place. We have investigated the number and type of microstates across this developmental window in two large geographically and culturally diverse cohorts from Cape Town, South Africa (Khula, N=318), and São Paulo, Brazil (Germina, N=536). We found consistent microstate classes emerged between the two disparate cohorts over development. For microstate classes that were present throughout the first two years, we also evaluated developmental changes in their durations and occurrences to understand largescale network dynamics in this period. We found both consistent and site-unique developmental patterns across microstate dynamics, suggesting these indices of largescale networks reflect both foundational and context-sensitive neurodevelopmental dynamics. These results inform our understanding about a key period of brain development in the following ways.

First, we found highly conserved patterns of microstate classes across two geographically and culturally diverse cohorts using data-driven clustering. Specifically, our approach for obtaining microstate maps suggested the presence of microstates A, B, C, D and E in the first year of life with the emergence of two new microstates F and G (and the absence of microstate C) in the second year of life. Notably, microstate C was identified only in early infancy in both cohorts and may reflect incomplete spatiotemporal segregation and integration of networks in this period. Relatedly, in adult microstate literature where the microstate labelling has been restricted to only the four canonical maps A-D, other microstates are merged and labeled Microstate C, especially Microstate F because of the spatial correlation as C and F share regions from the Default Mode Network,and Microstate E because of shared topographical characteristics despite functional differences (Bréchet et al., 2019; Tarailis et al., 2021; Tarailis et al., 2023; Khanna et al., 2014; Pascual-Marqui et al., 2014; Santarnecchi et al., 2017; Zanesco, 2024). Indeed, microstate C re-emerged if we forced a 4-microstate class solution within these developmental data (not reported). Here the consistent developmental shift from Microstate C to F and the instability of microstate C in a data-driven classification context suggests incomplete integration of the default mode network in infancy. However, the overall pattern of microstate classifications suggests consistent largescale network configurations and developmental shifts in these configurations are observable in early life across disparate contexts.

Next, we found conserved evidence that faster transitions between largescale brain topographies emerge over infancy, especially over the first year of life. That is, the duration of all microstates (A, B, D, and E) significantly decreased within individuals over the first two years of life in both cohorts. Moreover, the best-fitting models for these developmental changes overwhelmingly included nonlinear terms (all except microstate A in Khula and microstate D in Germina), indicating the steepest decreases in the durations of microstates occurred over early infancy across both cohorts. These reductions in durations, reflecting faster transitions between largescale brain topographies, suggest more efficient sub-second level interactions between underlying largescale network configurations over infancy (Khanna et al., 2015; Michel & Koenig, 2018). Such developmental trajectories are consistent with fMRI findings in sleep indicating that infants undergo rapid increases in functional segregation and integration across brain regions, contributing to more efficient neural processing in largescale networks on the scale of seconds to minutes (Doria et al., 2010; Gao, Alcauter, Smith, et al., 2015; Gao et al., 2009; Lin et al., 2008). Moreover, the white matter tracts involved in sensory and cognitive networks undergo substantial maturation and myelination over the first two years of life to support faster conduction speed of neural signals and more efficient neural communication across distant brain regions (Deoni et al., 2016; Lebel & Deoni, 2018; Dubois et al., 2014; Gao et al., 2015b). These significant structural changes potentially facilitate the significantly shorter microstate durations that we observed in our results. Future research should examine how these early structural and functional indices of largescale networks co-develop.

Furthermore, sensory microstates showed consistent patterns of development between cohorts. Specifically, microstates A and B, associated with the auditory and visual network, respectively, became briefer with age, but recurred more frequently within each second. This temporal pattern aligns with existing research indicating that sensory networks emerge early postnatally and are among the first to mature during infancy (Michel & Koenig, 2018; Fair et al., 2009; Gao et al., 2015; Doria et al., 2010; Smyser et al., 2011). Here we show that this functional maturation in awake behaving infants includes rapid, frequent instances of these sensory network configurations each second, which may aid in the detection of relevant sensory information from infants’ environments. How these networks’ frequent, brief configurations at the sub-second level support learning and higher-order neurodevelopment should be explored further (Hill et al., 2023; Britz et al., 2010).

In contrast, the attention and task switching related microstates, Microstates D (dorsal attention network) and E (salience network), exhibited divergent developmental trajectories between the Khula and Germina cohorts. The dorsal attention network is crucial for voluntary attentional control, while the salience network is involved in detecting and integrating stimuli that are emotionally or behaviorally relevant (Menon & Uddin, 2010; Seeley et al., 2007; Zhou et al., 2018). Microstate D increased in occurrence with age in the Khula cohort but decreased in the Germina cohort, while Microstate E displayed the opposite pattern. The contrasting trajectories may reflect environmental or socio-cultural factors that shape higher-order brain network engagement in different ways across contexts. For example, prior research has suggested that differences in early caregiving environments, linguistic exposure, and socio-economic conditions can influence the pace of brain network maturation and integration (Tarailis et al., 2021; Bagdasarov et al., 2022; Khazaei et al., 2021; Holz et al., 2023). This pattern of findings highlights the need for more culturally and geographically diverse studies investigating neurodevelopment to unravel how geocultural conditions shape brain network organization and dynamics in early life.

In addition to these developmental changes, our results revealed small yet significant sex-related differences in several microstates’ dynamics. Compared to females, males across both cohorts showed significantly longer durations of microstate B, microstate D, and microstate E (in *Khula* only). We also observed significantly fewer occurrences of microstates A (*Khula* and *Germina*) and B (*Khula* only) in males as compared to females. Overall, males exhibited slightly longer durations and fewer occurrences of largely sensory microstates as compared to females, indicating slower temporal transitions between brain network states over infancy. This set of findings is consistent with prior studies indicating sex differences in neural maturation rates, particularly in brain regions associated with attention and sensory processing where males can lag females developmentally (Bagdasarov et al., 2022; Tomescu et al., 2018). Together, these different dynamics suggest potential sex differences in how largescale networks supporting sensory processing are engaged functionally at the sub-second level during early neurodevelopment. Such differences may have implications for understanding sex-specific developmental trajectories in cognitive domains, as slower transitions between neural states have been linked to later development of cognitive flexibility and executive function (Das et al., 2022; Hill et al., 2023). Longitudinal studies beyond age two years may shed light on how these foundational sex-dependent largescale network differences scaffold higher-order neurocognitive development.

### Limitations and future directions

While our study provides important insights into the developmental trajectories of EEG microstates across the first two years of life, several limitations and avenues for future research warrant consideration. First, our investigation is limited to the age range of 2 to 30 months. Extending studies beyond this range is crucial to capture the continued development of large-scale brain networks and their relationship to emerging cognitive and behavioral skills in early childhood. Additionally, while we examined microstate dynamics across two geoculturally diverse cohorts, future studies should include data from additional geographic, cultural, and socioeconomic contexts to further test the generalizability of microstate developmental patterns. Within cohorts, there is also a need to explore differences in brain network development as indexed by microstates as a function of psychosocial and environmental factors. While EEG microstates offer a temporally precise measure of brain dynamics, their spatial resolution remains limited. Future research could integrate EEG with other neuroimaging modalities like fMRI, to uncover the spatial correlates of microstate dynamics and to provide a more comprehensive understanding of early brain network organization (Nishida et al., 2013; Rajkumar et al., 2021). Moreover, our study focused on resting-state EEG, which limits knowledge about network dynamics in active cognition. Exploring task-related microstate dynamics in awake, behaving infants presents an exciting opportunity to elucidate how these networks support emerging cognitive and behavioral skills during development. Such integrative and multimodal approaches will help to advance our understanding of the temporal, spatial, and functional aspects of large-scale brain network maturation in early life.

### Conclusion

Our results illustrate both the early emergence and significant developmental changes in largescale brain networks’ functional dynamics at the sub-second level across infancy. By examining microstate properties across longitudinal cohorts diverse in geography, culture, and socioeconomic factors, our study underscores the robustness of EEG microstates as markers of early foundational brain development. Microstate trajectories across cohorts suggest some foundational sensory network functional development proceeds similarly across these geocultural differences while networks serving higher-order cognitive processes like attention are sensitive to contextual differences. Together these longitudinal results provide new insights into how largescale functional brain network dynamics and development unfold in early life.

## Methods

### Participants

#### Cohort 1: Khula

394 mothers and their infants were recruited for this study (out of which 329 were enrolled antenatally) from Gugulethu, an informal settlement in Cape Town, South Africa, as part of a larger longitudinal project (Zieff et al., 2024). Women were eligible to participate in the study if they were (i) in their third trimester of pregnancy (28-36 weeks) or up to three months postpartum, and (ii) over the age of 18 years at the time of recruitment. Inclusion criteria for the study included (a) singleton pregnancy, (b) no psychotropic drug endorsement during pregnancy, (c) no infant congenital malformation or abnormalities (e.g., spina bifida, Down’s syndrome), (d) no significant delivery complications (e.g., uterine rupture, birth asphyxia), and e) gestational age 36 weeks or greater. Study demographics are provided in **Table 1**. *In Xhosa language, Khula means “to grow”, signifying development*.

Families were invited to participate in four in-lab study visits with EEG over their infant’s first two years of life. Not all infants contributed usable EEG at every visit. 11.3% infants provided usable EEG data at only 1 timepoint, 13.5% infants provided 2 timepoints, 34% infants provided 3 timepoints, and 41.2% infants provided data at all 4 timepoints. The first visit occurred when infants were between approximately 2 months and 6 months of age (mean age from usable data: 3.74 months; 51.7% males), the second visit when they were between 5 and 12 months (mean age: 8.72 months; 50.6% males), the third visit when they were between 12 and 18 months (mean age: 14.1 months; 51.7% males) and finally, the fourth visit when they were between 19 and 26 months of age (mean age: 21.5 months; 51.8% males). 318 of the 394 enrolled infants contributed EEG data to the present study.

All procedures were approved by the Human Research Ethics Committee at the University of Cape Town in South Africa. Written consent was obtained from caregivers annually, giving caregivers more agency to opt out if they chose to discontinue participation at any point. Families were compensated with ZAR 300 (∼ USD 15) for their time after each visit. Additionally, transportation was made available to and from research sites for participants’ convenience. Food and beverages were also provided during each visit.

#### Cohort 2: Germina

557 mother-infant dyads were enrolled into a longitudinal cohort study in São Paulo, Brazil (for more information, see Fatori et al., 2024). Infants included in this analysis were recruited postnatally with the average age at recruitment being 2.4 months. Inclusion criteria were (a) maternal age of 20-45 years, (b) infant age of 3 months 0 days to 3 months and 29 days for the first visit, (c) gestational age of >37 weeks, (d) infant birth weight >2000 grams, (e) no substance endorsement during pregnancy, (f) no history of severe maternal mental disorders (e.g., psychosis, bipolar disorder), (g) no delivery complications requiring medical intervention (e.g. perinatal asphyxia, shoulder dystocia, excessive bleeding), (h) no infant genetic syndrome or auditory/visual impairment diagnoses, and (i) availability for in-person lab visits. Study demographics are provided in **Table 2***. In Portuguese, Germina means “to germinate”, signifying early growth*.

Families were invited to participate in four in-lab study visits with EEG over their infant’s first two years of life. Not all infants contributed usable EEG at every visit. 15.1% infants provided usable EEG data at only 1 timepoint, 33% infants provided 2 timepoints, 37.9% infants provided 3 timepoints, and 14% infants provided data at all 4 timepoints. The first visit occurred when infants were between approximately 3 months and 4 months of age (mean age from usable data: 3.62 months; 45.3% males), the second visit when they were between 5 and 10 months (mean age: 7.64 months; 48.9% males), the third visit when they were between 10 and 17 months (mean age: 13.2 months; 48.9% males) and the fourth visit when they were between 18 and 30 months of age (mean age: 19.7 months; 50.2% males). EEG data from 536 of the 557 enrolled infants were deemed usable for the study.

All procedures were approved by the Ethics Committee for the Analysis of Research Projects (CAPPESq) and the National Council of Ethics in Research (ref.: CAAE 49671221.2.0000.0068). In accordance with the Declaration of Helsinki, all mothers provided written informed consent before completing any study measure.

### EEG data acquisition

Baseline EEG was recorded at each visit using largely harmonized a priori protocols across visit ages and between sites. Infants were seated on their caregiver’s lap approximately 60 cm in front of a computer monitor (30 x 45.5 cm, 1440 x 900-pixel resolution) throughout recording where they passively viewed a video of moving shapes as in (Fox et al., 2024). The video was programmed and presented in E-Prime (Psychology Software Tools, Inc., Sharpsburg, PA). Recordings took place in a dimly lit, quiet room without electrical shielding. EEG was recorded using a 128-channel HydroCel geodesic sensor net and a Net Amps 400 amplifier (Magstim EGI, Whitland, UK). Data were referenced online to the vertex electrode Cz and sampled at 500Hz (Germina) and 1000Hz (Khula) via NetStation software (Magstim EGI, Whitland, UK). Electrode impedances were kept below 100 kΩ wherever possible in accordance with the capabilities of the high-impedance amplifiers. In Khula, nets designed with modified taller pedestals were used as needed for improving the inclusion and experience of infants with Afro-textured hair (Mlandu et al., 2024). Shea moisture leave-in castor oil conditioner was applied to hair across the scalp prior to net placement to improve both impedances and participant comfort. The leave-in conditioner is insulating to prevent electrical bridging, has not been found to disrupt the EEG signal during testing, and allows for nets to lay closer to the scalp for Afro-textured hair types, while making it far more comfortable to remove from the scalp at the end of testing (Fox et al., 2024; Mlandu et al., 2024).

### EEG data preprocessing

EEG data were converted from native Netstation .mff format to .raw format to remove identifying video information across sites. The .raw files were processed using the Harvard Automated Processing Pipeline for EEG (HAPPE), an automated open-source EEG preprocessing software customized for the developmental population (Gabard-Durnam et al., 2018; Monachino et al., 2022). HAPPE v 4.1 was run using MATLAB (2022b) and EEGLAB (Delorme & Makeig, 2004) with the preprocessing parameters as listed in Supplementary Table 1. As commonly used for microstate analysis, the data were filtered between 1-40 Hz (Britz et al., 2010) with a finite impulse response (FIR) bandpass filter in HAPPE. The outer rim electrodes of the net were removed from analysis, which is a common practice in infant EEG research given their susceptibility to artifact contamination (see Monachino et al., 2022). Bad channel detection was performed on remaining channels. Electrical line noise was removed at 50 Hz from the *Khula* data and at 60 Hz from the *Germina* data using CleanLine (Mullen, 2012) via a multi-taper regression which can remove electrical noise without distortion of the EEG signal in the nearby frequencies, unlike traditional notch filters (Mitra & Pesaran, 1999). Artifacts were corrected with primarily wavelet-thresholding, but also MuscIL (independent component analysis restricted to muscle-classified artifacts) within HAPPE. Bad channels were interpolated followed by rereferencing of the data to the average reference. The continuous EEG recording from each participant was epoched into 2-second-long segments and any segment with amplitude change ±150µV was further rejected. See Supplementary Table 2 for EEG data quality measures across sites.

### Microstate computations

Resting state EEG microstates in infants were computed from the preprocessed EEG data using the *generateMicroStates.m* script of HAPPE v4.1, which is based on functions from EEGLAB (Delorme & Makeig, 2004) and the microstate toolbox plugin (MST version 1.0) (Poulsen et al., 2018). Participants who retained more than 15 segments (i.e., 30 seconds of data) after preprocessing were considered for further analysis. Given the paucity of literature on pre-processing for microstates in early development (though see Bagdasarov et al., 2024), additional sensitivity analyses were conducted to test whether the identity and number of microstate classes identified in the data varied as a function of retained segments by re-computing microstates using only infants with 100% retained segments (91 segments in at least 30 infants). These restricted analyses returned the same number and type of microstate classes as reported with the full sample, suggesting microstate determination was robust to segment retention variance within this developmental sample and consistent with prior research on the matter (Bagdasarov et al., 2024).

For each cohort, microstate analysis was performed for each visit (timepoint) separately to determine whether identity and number of microstate classes varied over early development. To identify microstate classes, all available segments from a participant were concatenated and treated as one continuous signal. Datapoints corresponding to GFP (Global Field Power) maxima, which represent maps with a high signal-to-noise ratio, were subjected to a modified k-means clustering algorithm (Koenig et al., 2002; Pascual-Marqui et al., 1995) to identify microstate prototypes (within the range of 2 to 8 prototypes) in a polarity invariant manner. This means that the algorithm did not differentiate between spatially proportional but oppositely polarized topographical maps when assigning microstate clusters (Pascual-Marqui et al., 1995). The assumption in clustering is that all EEG data assigned to the same cluster originate from neural generators underlying the prototype topography of that cluster. The clustering process was initialized 50 times stochastically, with a maximum of 1,000 iterations per run. A total of 1,000 GFP peaks per participant were included in the segmentation, with a minimum peak distance of 10 ms, as recommended by Poulsen et al. (2018). Visual inspection of the topographies and evaluation of fit measures, including global explained variance (GEV) and the cross-validation (CV) criterion (Poulsen et al., 2018), revealed 5 to 6 prototypical microstate maps (classes A to G) that provided an optimal clustering solution depending on the ages being clustered. While higher GEV indicated better explained variance, a lower CV criterion reflected less residual noise and was thus desirable. This data-driven selection represented a qualitative balance between specificity and generalizability across cohorts and timepoints, where the former typically improved by increasing the number of microstates and the latter by limiting their number (Hill et al., 2023; Michel & Koenig, 2018). The microstate classes obtained were subsequently backfitted to each participant’s EEG data. To mitigate the influence of noise inherent in spontaneous EEG recordings, which can result in consecutive short time frames being labeled as different classes (Musaeus et al., 2019; Poulsen et al., 2018), we applied the default temporal smoothing approach in the microstate toolbox and rejected microstate segments shorter than 30 ms (Poulsen et al., 2018). This method reclassifies the labels of those segments to the next best-fitting microstate, as measured by global map dissimilarity (Poulsen et al., 2018). The following statistical parameters were then extracted from the microstate labeled data corresponding to the various microstate classes at each visit: (a) *Duration,* defined as the average duration in milliseconds for which a given microstate class remains stable; (b) *Occurrence* determines the average number of times a microstate class is dominant within one second; (c) *Coverage* reflects the fraction of time for which a given microstate is active uninterruptedly; and (d) *GEV* or the global explained variance, is the variance in the EEG data explained by a particular microstate class.

### Statistical Analyses

The mean and the standard deviation of each microstate parameter (i.e. occurrence, duration, coverage, GEV) were computed across participants within each cohort for each of the microstate classes for all four visits. Datapoints lying outside 3 standard deviations from the mean for a given microstate feature for each visit within cohort were considered outliers and were not included in the analyses. Using this criterion, 11 datapoints were removed from Khula analyses (1 at *timepoint 1*, 3 at *timepoint 2*, none at *timepoint 3*, and 7 at *timepoint 4*) and 6 datapoints were removed from Germina analyses (none at *timepoint 1*, 1 at *timepoint 2*, 2 at *timepoint 3*, and 3 at *timepoint 4*).

Primary statistical analyses focused on duration and occurrence features for the microstate classes present across all 4 visits in each cohort. For each cohort, the developmental changes in the durations and the occurrences of the microstate classes were analyzed with linear mixed effect models using the *lme4* package in *R version 2024.04.2+764* (R Core Team, 2024). The model parameters were estimated using maximum likelihood (Bates et al., 2015). Random subject intercepts were included in all models to account for between-participant variability in initial microstate characteristics. A fixed effect of sex was also modeled, and the number of retained segments at each visit was included as a covariate of no interest given prior research suggesting some microstate features’ internal consistency varies with segment number (Bagdasarov et al., 2024). Retained segment number had a negligible effect size in all models (η_p_^2^ < 0.01).

To assess the best-fitting model for the age-associated growth trajectories of microstate features within each cohort, three models of age-related changes were run for each feature that included a) linear age term, b) linear and quadratic age term, c) linear and logarithmic age term. The fits of these three models were compared (*linear* vs *linear+quadratic* vs *linear+log*) using BIC (Bayesian Information Criterion) to reduce the risk of over-fitting data by penalizing the more complicated models for additional terms. A nonlinear, more complex model was selected if there was strong support for inclusion of the nonlinear term (i.e. BIC difference of 6 points or greater), and if both nonlinear term models (*linear+quadratic* vs *linear+log*) met the BIC criterion for support, the model with the lowest BIC was chosen as the best fit to the data (Raftery, 1995).

## Funding

This research has been supported by the Wellcome LEAP 1kD Program (KD, GVP, ES, LGD).

## Competing interests

The authors declare no competing interests.

---

**Supplementary Table 1.** HAPPE v4 Preprocessing Script Parameters.

| ***Preprocessing Parameter Type*** | ***Parameter Used*** |
| --- | --- |
| **Density** | High (>30 channels) |
| **Resting State or Task** | Resting state |
| **Acquisition Layout** | 128 channel EGI HydroCel Geodesic Sensor Net |
| **Channels** | All except E1, E8, E14, E17, E21, E25, E32, E38, E43, E44, E48, E49, E56, E68, E73, E81, E88, E94, E99, E107, E113, E114, E119, E120, E121, E125, E126, E127, E128 |
| **Line Noise Frequency** | 50 Hz (South Africa) / 60 Hz (Brazil) |
| **Line Noise Reduction Method** | CleanLine - Default |
| **Resample** | Off |
| **Filter - Lowpass Cutoff** | 40 Hz |
| **Filter - Highpass Cutoff** | 1 Hz |
| **Filter Type** | EEGLAB's FIR |
| **Bad Channel Detection** | On |
| **Bad Channel Detection Method** | Default |
| **Wavelet Thresholding** | Default |
| **Wavelet Threshold Rule** | Hard |
| **MusciL** | On |
| **Segmentation** | On |
| **Segment Length** | 2 seconds |
| **Interpolation** | Off |
| **Segment Rejection** | On |
| **Segment Rejection Method** | Amplitude criteria only |
| **Minimum Segment Rejection Threshold** | -150 μV |
| **Maximum Segment Rejection Threshold** | 150 μV |
| **Re-Reference Method** | Average |

**Supplementary Table 2.** EEG Quality Control Measures.

| *Cohort* |  | *Khula* |  |  |  | *Germina* |  |  |  |
| --- | --- | --- | --- | --- | --- | --- | --- | --- | --- |
| EEG quality control measure |  | Visit 1 (N=242) | Visit 2 (N=249) | Visit 3 (N=261) | Visit 4 (N=218) | Visit 1 (N=349) | Visit 2 (N=444) | Visit 3 (N=350) | Visit 4 (N=201) |
| % channels selected |  | 89.72 (6.9);<br>57.58 - 100 | 93.75 (4.84);<br>71.72 - 100 | 92.63 (5.3);<br>64.65 - 100 | 91.83 (6.3);<br>64.65 - 100 | 85.43 (6.9);<br>41 - 97 | 89.58 (5.21);<br>61 - 98 | 90.22 (4.25);<br>76 - 99 | 90.8 (4.89);<br>68 - 99 |
| % trials retained |  | 94.34 (15.46);<br>16.48 - 100 | 94.86 (15.2);<br>16.48 - 100 | 95.87 (10.71);<br>26.37 - 100 | 96.33 (12.5);<br>23.08 - 100 | 93.81 (15.34);<br>23.08 - 100 | 90.06 (18.33);<br>24.19 - 100 | 89.35 (19.3);<br>24.59 - 100 | 93.44 (15.63);<br>28.33 - 100 |
| Correlation of data pre- vs post- wavelet thresholding (Pearson's r) | at 2 Hz | 0.41 (0.2);<br>0.00005 - 0.87 | 0.34 (0.21);<br>0.001 - 0.84 | 0.38 (0.22);<br>0.003 - 0.9 | 0.48 (0.22);<br>0.01 - 0.95 | 0.33 (0.22);<br>0.01 - 0.9 | 0.35 (0.24);<br>0.0001 - 0.91 | 0.32 (0.25);<br>0.003 - 0.92 | 0.37 (0.24);<br>0.003 - 0.96 |
|  | at 5 Hz | 0.63 (0.23);<br>0.0002 - 0.99 | 0.61 (0.27);<br>0.004 - 0.95 | 0.64 (0.25);<br>0.02 - 0.97 | 0.76 (0.21);<br>0.02 - 0.99 | 0.52 (0.26);<br>0.01 - 0.96 | 0.49 (0.27);<br>0.0009 - 0.95 | 0.48 (0.29);<br>0.01 - 0.97 | 0.54 (0.28);<br>0.01 - 0.98 |
|  | at 8 Hz | 0.6 (0.24);<br>0.0003 - 0.98 | 0.64 (0.26);<br>0.003 - 0.98 | 0.7 (0.26);<br>0.03 - 0.98 | 0.83 (0.18);<br>0.02 - 0.99 | 0.46 (0.26);<br>0.02 - 0.91 | 0.49 (0.28);<br>0.0008 - 0.97 | 0.53 (0.3);<br>0.01 - 0.99 | 0.62 (0.29);<br>0.01 - 0.99 |
|  | at 12 Hz | 0.63 (0.24);<br>0.0007 - 0.98 | 0.59 (0.27);<br>0.01 - 0.95 | 0.63 (0.27);<br>0.01 - 0.97 | 0.77 (0.21);<br>0.01 - 0.99 | 0.5 (0.26);<br>0.01 - 0.94 | 0.5 (0.27);<br>0.0007 - 0.96 | 0.48 (0.29);<br>0.01 - 0.97 | 0.55 (0.27);<br>0.01 - 0.98 |
|  | at 20 Hz | 0.71 (0.21);<br>0.0002 - 0.99 | 0.65 (0.25);<br>0.01 - 0.96 | 0.67 (0.25);<br>0.01 - 0.97 | 0.78 (0.19);<br>0.01 - 0.99 | 0.6 (0.24);<br>0.04 - 0.95 | 0.59 (0.26);<br>0.0009 - 0.96 | 0.56 (0.27);<br>0.01 - 0.98 | 0.61 (0.26);<br>0.02 - 0.98 |
|  | at 30 Hz | 0.76 (0.2);<br>0.0001 - 0.99 | 0.7 (0.24);<br>0.01 - 0.97 | 0.71 (0.25);<br>0.01 - 0.98 | 0.79 (0.17);<br>0.03 - 0.99 | 0.67 (0.23);<br>0.04 - 0.97 | 0.67 (0.25);<br>0.0009 - 0.98 | 0.62 (0.26);<br>0.01 - 0.99 | 0.67 (0.25);<br>0.03 - 0.98 |

## Notes

### Competing Interest Statement

The authors have declared no competing interest.

